# CD8 T cell epitope generation toward the continually mutating SARS-CoV-2 spike protein in genetically diverse human population: Implications for disease control and prevention

**DOI:** 10.1101/2020.09.10.290841

**Authors:** Elisa Guo, Hailong Guo

**Affiliations:** Mounds View High School, 1900 Lake Valentine Rd, Arden Hills, MN 55112; Independent scientist, St Paul, MN 55126

## Abstract

The ongoing pandemic of SARS-CoV-2 has brought tremendous crisis on global health care systems and industrial operations that dramatically affect the economic and social life of numerous individuals worldwide. Understanding anti-SARS-CoV-2 immune responses in population with different genetic backgrounds and tracking the viral evolution are crucial for successful vaccine design. In this study, we reported the generation of CD8 T cell epitopes by a total of 80 alleles of three major class I HLAs using NetMHC 4.0 algorithm for the spike protein of SARS-CoV-2, a key antigen that is targeted by both B cells and T cells. We found diverse capacities of S protein specific epitope presentation by different HLA alleles with very limited number of predicted epitopes for HLA-B*2705, HLA-B*4402 and HLA-B*4403 and as high as 132 epitopes for HLA-A*6601. Our analysis of 1000 S protein sequences from field isolates collected globally over the past few months identified three recurrent point mutations including L5F, D614G and G1124V. Differential effects of these mutations on CD8 T cell epitope generation by corresponding HLA alleles were observed. Finally, our multiple alignment analysis indicated the absence of seasonal CoV induced cross-reactive CD8 T cells to drive these mutations. Our findings provided molecular explanations for the observation that individuals with certain HLA alleles such as B*44 are more prone to SARS-CoV-2 infection. Studying anti-S protein specific CD8 T cell immunity in diverse genetic background is critical for better control and prevention of the SARS-CoV-2 pandemic.

## Introduction

The coronavirus (CoV) is an enveloped, positive-stranded RNA virus that can cause respiratory and enteric diseases in wide range of hosts including human, numerous animals, birds and fish(1). Four genera (Alpha, Beta, Gamma, and Delta) of CoVs have been classified with human CoVs designated within Alpha and Beta groups. Their genome, about 30kb in length, is the largest found in RNA viruses and encodes more than 20 putative proteins, including four major structural proteins: spike (S), envelope (E), membrane (M), and nucleocapsid (N)(1). The human seasonal CoVs are endemic throughout the world, causing 15-30% respiratory tract infections that are typically mild and self-limiting. However, the outbreak of severe acute respiratory syndrome (SARS) in 2003(2) and Middle East respiratory syndrome (MERS) in 2012(3) that were caused by the infection of SARS-CoV and MERS-CoV had led to about a mortality rate of about 10% and 40%, respectively(1,4).

In December 2019, a novel coronavirus disease (COVID-19), caused by 2019 novel coronavirus (2019-nCoV) and later renamed as severe acute respiratory syndrome coronavirus 2 (SARS-CoV-2), was identified in Wuhan City, Hubei Province, China from patients with severe pneumonia(5–7). Subsequently, this novel viral infection has rapidly spread into nearly all countries over the world, leading to the declaration of the first-known coronavirus global pandemic by the World Health Organization (WHO) on March 11, 2020(8). As of July 31, the COVID-19 pandemic has resulted in over 17 million confirmed cases and more than 680,000 deaths globally according to the update from Johns Hopkins Coronavirus Resource Center (https://coronavirus.jhu.edu/map.html). Like SARS-CoV and MERS-CoV, SARS-CoV-2 was identified as betacoronavirus through genome sequencing and bioinformatic analysis(6,7) and has been formally classified by the International Committee on Taxonomy of Viruses (ITVC)(9).

The S protein of SARS-CoV contains neutralization epitopes, which are critical for eliciting anti-viral B cell immune responses. Therefore, S protein appears to be a promising antigen target for vaccine development(4,10). Similarly, a growing body of research has identified neutralization epitopes on SARS-CoV-2 S protein(11–13). Although SARS-CoV neutralization antibody is critical, the neutralizing antibody titers and memory B cell response were short-lived in SARS-recovered patients(14). Similarly, the antibodies including the neutralization antibody induced by SARS-CoV-2 infection could fade quickly(15,16). In addition to the anti-S protein humoral response, S protein specific CD8 T cells have been shown with protective immunity during SAR-CoV infection and are critical for vaccine efficacy(17,18). Therefore, identifying SARS-CoV-2 S protein specific CD8 T cell epitopes and fully utilizing CD8 T cell immunity in addition to neutralization response are urgently needed for studying anti-COVID-19 immunity and developing effective vaccines. Since its initial outbreak, SARS-CoV-2 has undergone genetic mutations(19). Further, mutations on S proteins that may affect human ACE2 receptor binding, viral transmission and infectivity have been reported(20–22). However, the potential effects of these mutations on CD8 T cell epitope generation have not been analyzed.

In this study, we used the NetMHC 4.0 prediction algorithm to analyze a panel of classical human MHC I (HLA-A, B and C) restricted CD8 epitopes on the S protein of SARS-CoV-2. Our data showed a significant variation of CD8 T cell epitope repertoires for different HLA-A, HLA-B and HLA-C alleles. In addition, our analysis using a large number of S protein sequences derived from SARS-CoV-2 genomes revealed important mutations including L5F, D614G and G1124V that are although less likely driven by seasonal CoV derived cross-reactive CD8 T cells, may differentially impact CD8 T cell epitope generation by different HLA alleles. Our results indicate genetic variability of three major class I HLAs may regulate S protein specific CD8 T cell responses, leading to differential susceptibility to and severity of the infection of SARS-CoV-2 that keeps mutating.

## Results

### S protein specific CD8 T cell epitopes of different HLA alleles

The NetMHC 4.0 algorithm has been widely utilized for predicting CD8 T cell epitopes of human and animals. By using this tool, our analysis for the S protein of reference Wuhan-Hu-1 virus generated a total of 5044 CD8 T cell epitope pool for three major class I HLAs with a total of 80 different alleles (Supplementary Table 2). Among these, there are 2282 epitopes for HLA-A, 1816 for HLA-B and 946 for HLA-C (Table 1). Within HLA-A, A*6601 allele showed 132 epitopes, the highest among all, indicating a potential broad and strong CD8 T cell responses against SARS-CoV-2 from individuals bearing this particular allele. However, there were several HLA-A alleles (*0201, A*0205, A*0207, A*0301, A*3101, A*3301) that only had about 30 epitopes. Other HLA-A alleles showed epitopes ranging from 40 to around 100 (Table 1). Similarly, within HLA-B, the total number of epitopes for individual alleles differed substantially, ranging from 16 to 118. The lowest number of epitopes among all the alleles analyzed was generated by HLA-B*2705. Additionally, HLA-B*4402 and HLA-B*4403 also had very low numbers of CD8 T cell epitopes (17 and 19 each) (Table 1). This observation may explain why B*44 positive individuals are more prone to SARS-CoV-2 infection(23). Furthermore, the total number of epitopes the HLA-C alleles could present varied from 75 to 121. These results indicate the capacity to present SARS-CoV-2 S protein specific CD8 T cell epitopes among different HLA alleles could be of great difference.

**Table 1.**
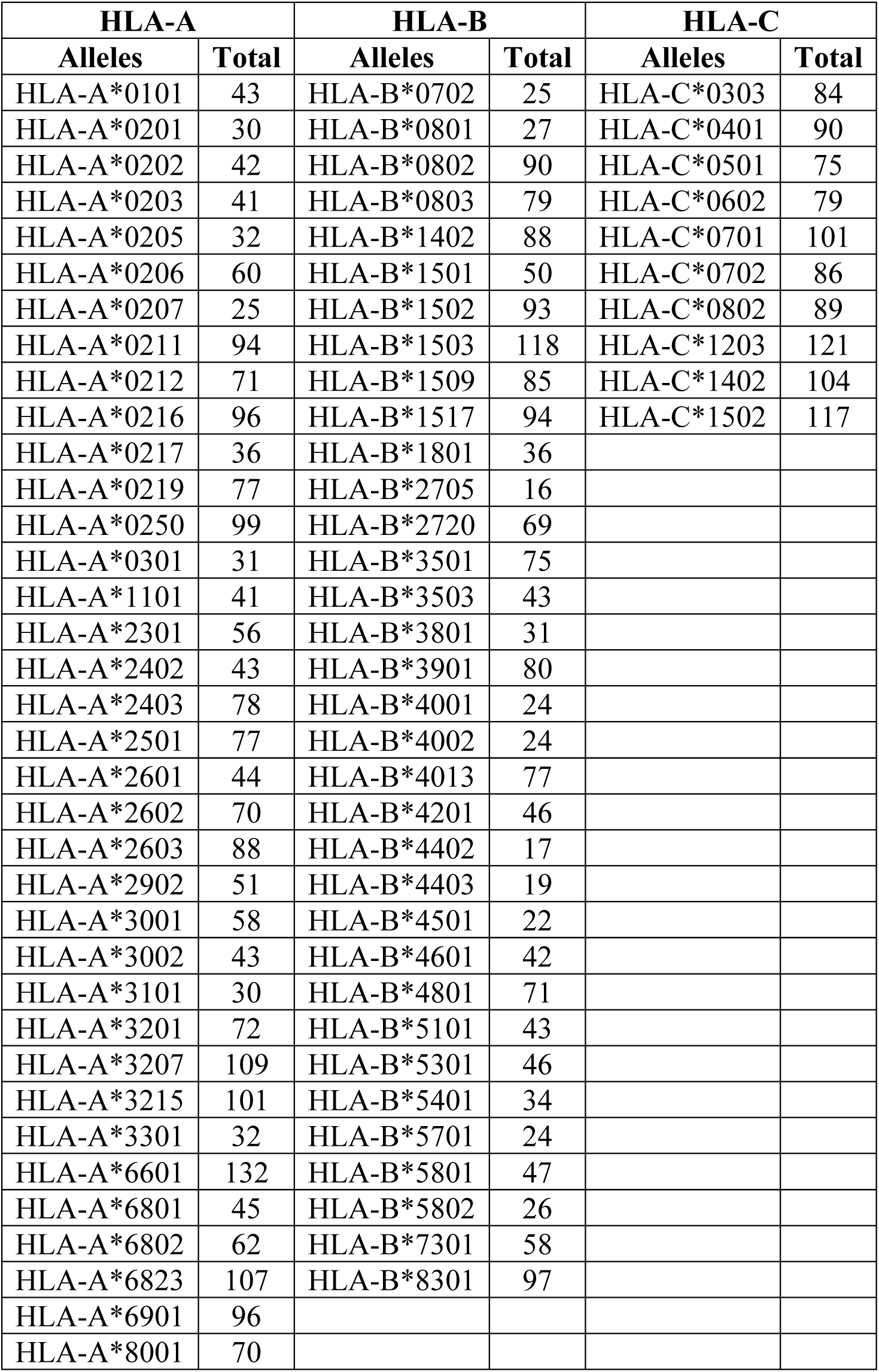
Summary of the total number of predicted SARS-CoV-2 S protein specific CD8 T cell epitopes for each HLA alleles

### Mutations on S protein of SARS-CoV-2

SARS-CoV-2 has evolved since its initial outbreak by changing its viral genome, which may cause mutations in viral proteins including S protein, resulting in the establishment of better fitness(19,24). We analyzed 1000 full length or near-full length S protein sequences (Supplementary Table 1) through multiple alignment and comparison with the reference S protein sequence to identify potential recurrent mutations and assess the effect on CD8 T cell epitope generation for different HLAs. The result in Fig 1 showed the location map of massive mutations across entire S protein that were observed from the 1000 S protein sequences. It appears that the hot spot targeted for mutation is located on the S1 subunit composed of the signal peptide region (Position 1-13), N terminal domain, NTD (Position 16-317), receptor binding domain, RBD (Position 330-521) and the subsequent SD1 and SD2 domains (Position 522-680) that is ahead of the S1/S2 cleavage motif PRRAR(25). Although there are plenty of positions where the mutation can occur, the frequency of these mutations is usually 0.1-0.2%, except three amino acids at position 5, 614 and 1124 that showed a frequency of 1.6%, 63% and 1.3%, respectively (Fig 1 and Supplementary Fig 1). These mutation rates are all beyond the preferred 0.5% mutation threshold that is used in influenza research(26) and the 0.3% tracking threshold for SARS-CoV-2 surveillance(27). In addition, these mutations are exclusively L to F at position 5, D to G at position 614 and G to V at position 1124. A single mutation on CD8 T cell epitope that is recognized and bound by HLA is one of the tactics several viruses, such as HIV, HCV and Influenza can use to escape host immunity(28–30). Therefore, we further analyzed the potential impact of these individual mutations on CD8 T cell epitope generation by different HLA alleles.

**Fig 1.**
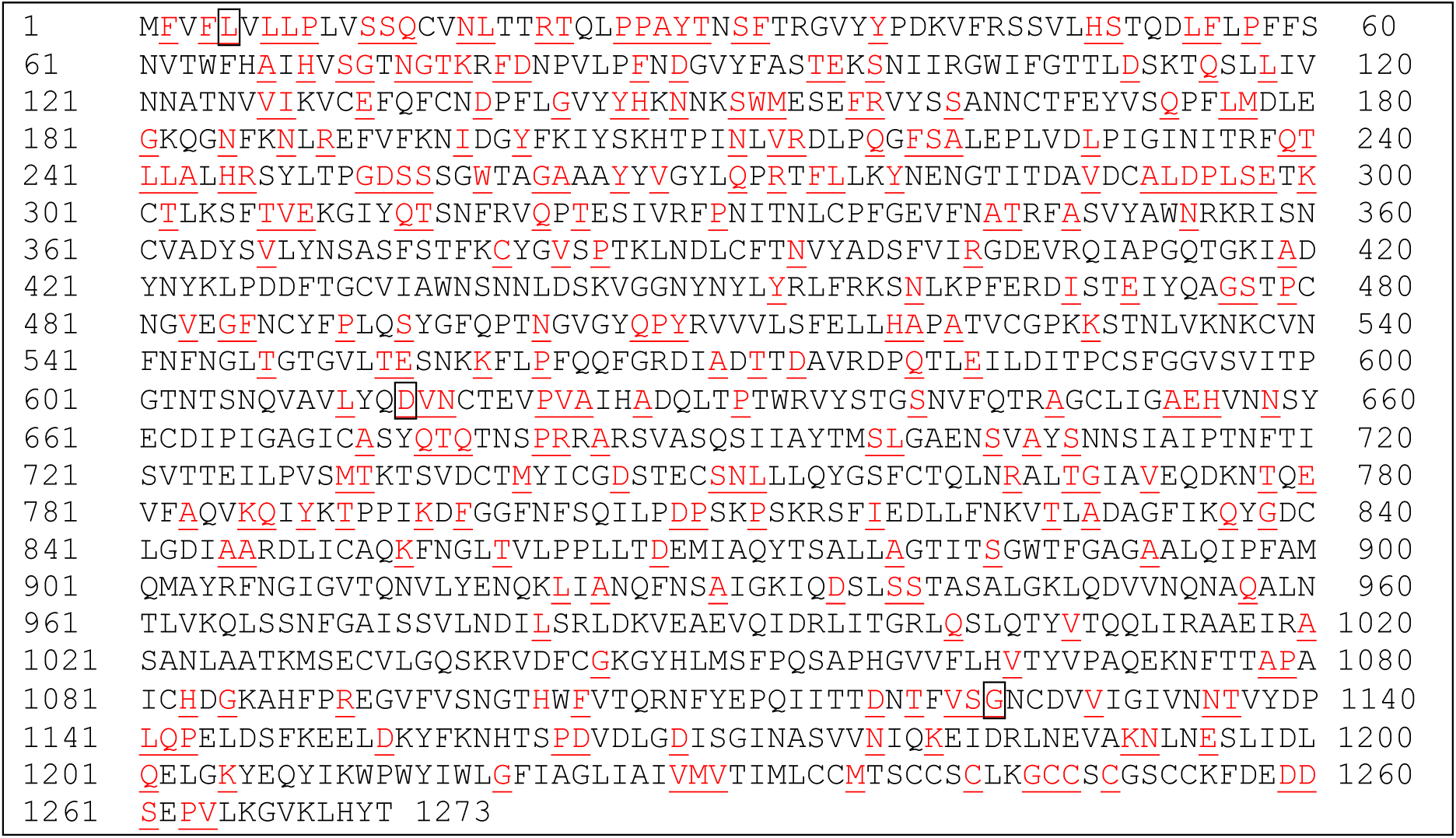
Distribution of mutations (in red and underlined) over the entire S proteins derived from multiple alignment analysis of 1000 S protein sequences of SARS-CoV-2 with the reference S protein as described. Mutation of the boxed amino acid at position 5 (L to F) was observed 16 times, at position 614 (D to G) 632 times and at position 1124 (G to V) 13 times, respectfully. All other mutations occurred less than 10 times, typically 1-2 times.

### Effect of L5F mutation on CD8 T cell epitope generation

The amino acid L on position 5 is part of the S protein signal peptide(25). Signal peptides are important targets of CD8 T cell responses against viral and tumor antigens(31,32). From the list of CD8 T cell epitopes for the 80 class I HLA alleles (Supplementary Table 2), we identified a total of 47 epitopes containing the L5 amino acid with the same sequence of FVFLVLLPL that could be presented by 22 of 36 HLA-As, 16 of 34 HLA-Bs and 9 of 10 HLA-Cs we analyzed (Table 2). Other potential epitopes containing the L5 are very limited and thus not evaluated. To examine whether a single mutation of L to F at position 5 on the SARS-CoV-2 S protein would affect FVFLVLLPL epitope presentation by different HLA alleles, we replaced the L with F at position 5 on the reference S protein sequence and re-analyzed the 9-mer CD8 epitopes for all the 80 HLA-A, B and C alleles using NetMHC 4.0 as described. The resulting epitopes of each allele were compared to the corresponding epitopes derived from the original reference S protein (Supplementary Table 2). The comparison data in Table 2 showed that L5F mutation increased the epitope binding affinity for 37 different HLA alleles, meanwhile only 10 other alleles had decreased binding affinity for the mutated epitope FVFFVLLPL. In addition, the mutated epitopes could be presented by 5 more HLA alleles (Table 2).

**Table 2.**
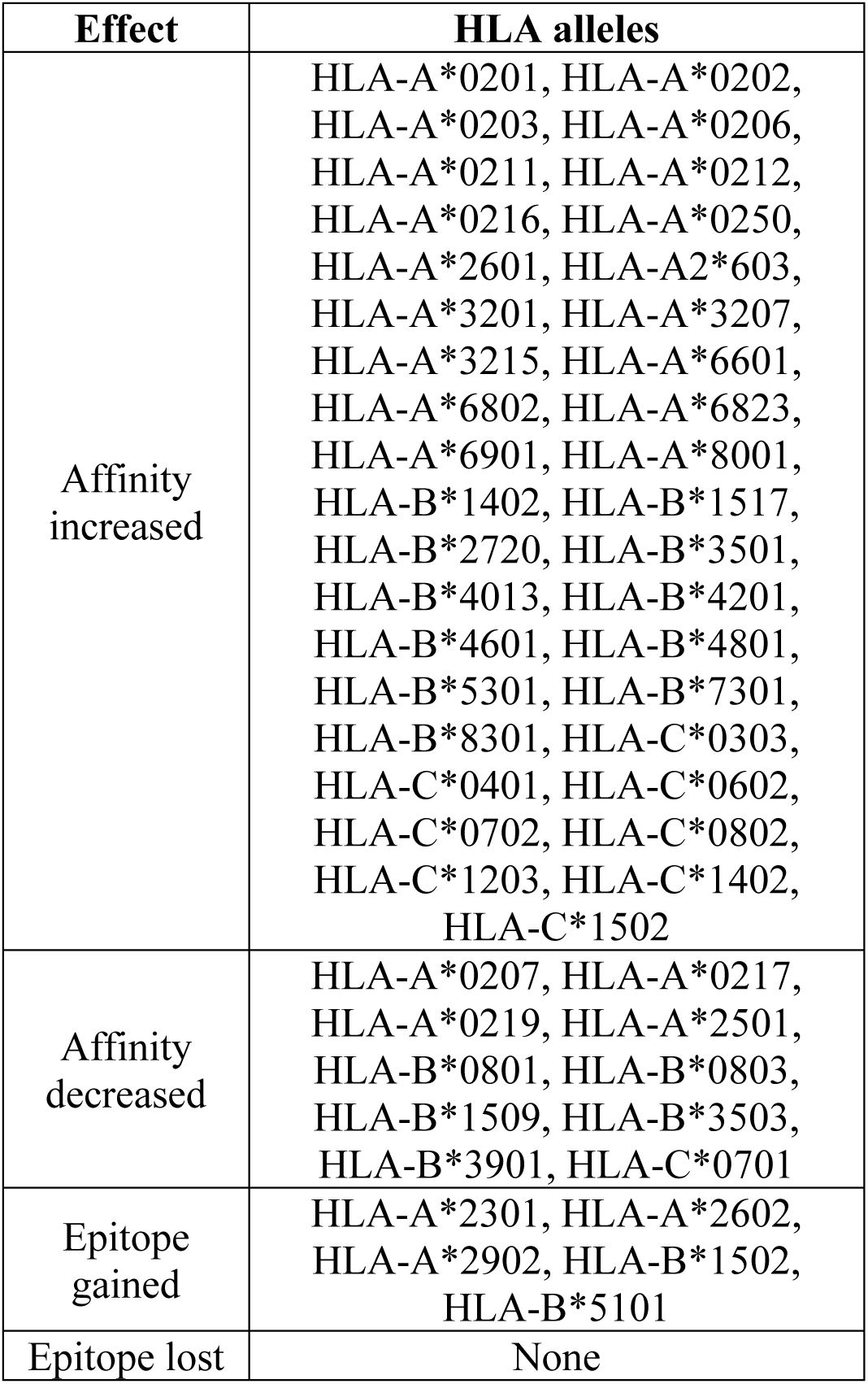
Effect of L5F mutation on the generation of SARS-CoV-2 S protein specific CD8 T cell epitope FVFFVLLPL by different HLA alleles

### Effect of G1124V mutation on CD8 T cell epitope generation

The amino acid G on position 1124 is within the connector domain of the S protein(25) that is important for S protein trimerization and critical for stabilizing S protein conformational structure during pre-and post-fusion with host cell membrane(33). By inspecting the epitopes in Supplementary Table 2, we found a total of 19 of the exact same epitopes starting at position 1121 (FVSGNCDVV) that could be presented by 12 HLA-As (HLA-A*0201, HLA-A*0202, HLA-A*0203, HLA-A*0206, HLA-A*0207, HLA-A*0211, HLA-A*0212, HLA-A*0216, HLA-A*0219, HLA-A*0250, HLA-A*6802 and HLA-A*6901) and 7 HLA-Cs (HLA-C*0303, HLA-C*0501, HLA-C*0602, HLA-C*0701, HLA-C*0802, HLA-C*1203 and HLA-C*1502). Only one additional epitope starting at a different position that contains the G1124 was identified for just one individual allele, indicating FVSGNCDVV is the predominantly presentable epitope in the connector domain of the S protein. Re-analyzing the epitopes of the 80 HLA alleles with the mutated reference S protein containing the V1124 identified 13 HLA alleles with decreased binding affinity for the mutated epitope FVSVNCDVV (Table 3). We also observed 6 other HLA alleles lost the ability to present the mutant epitope. No alleles showed increased binding affinity due to this mutation. Further there were no other alleles capable of presenting this mutated epitope (Table 3).

**Table 3.**
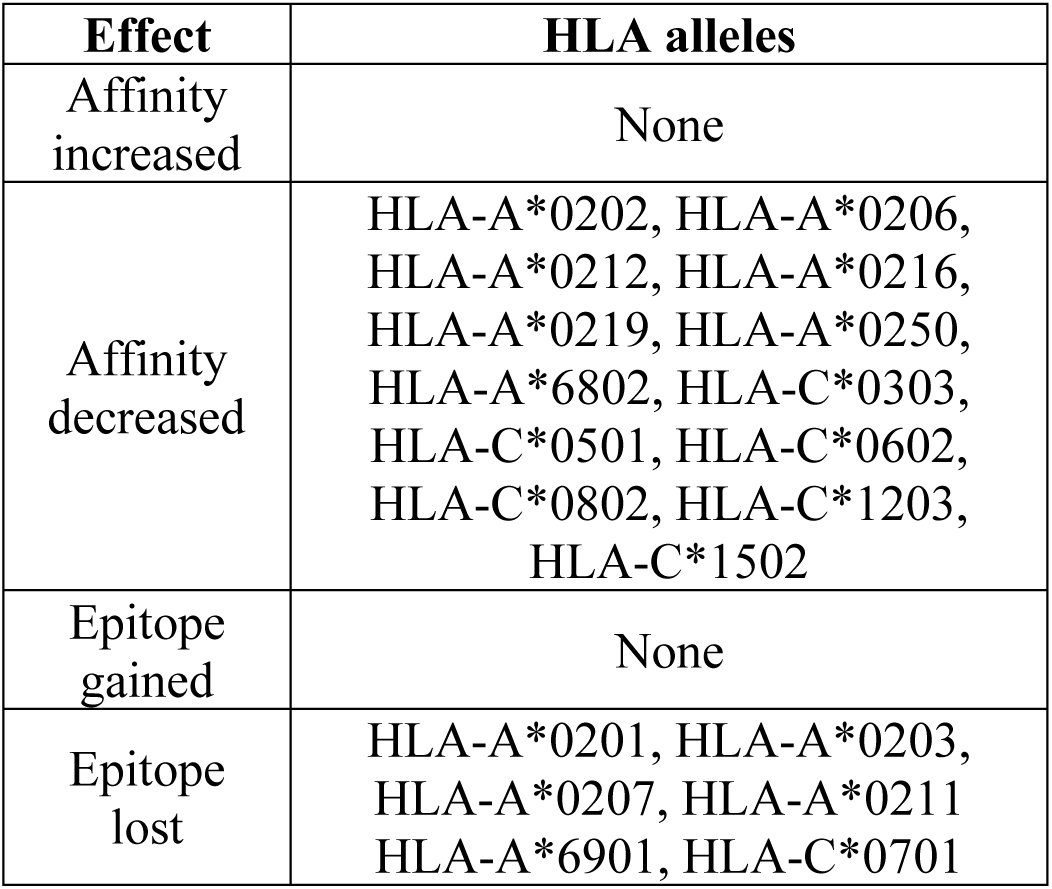
Effect of G1124V mutation on the generation of SARS-CoV-2 S protein specific CD8 T cell epitope FVSVNCDVV by different HLA alleles

### Effect of D614G mutation on CD8 T cell epitope generation

The mutation at position 614 with a sole D to G switch at a 63% frequency is especially alarming. The result revealed that SARS-Cov-2 isolates with G614 mutation has been adapted and spread efficiently within human population. To our knowledge, the D614 amino acid is not within the essential receptor binding domain or a residue of any validated neutralization epitopes for SARS-CoV-2. In addition, whether it is involved in CD8 T cell response is not known. Through screening the epitopes listed in Supplementary Table 2, we found that a panel of 22 HLA alleles including HLA-A*0101, HLA-A*0201, HLA-A*0206, HLA-A*0207, HLA-A*0211, HLA-A*0212, HLA-A*0216, HLA-A*0219, HLA-A*0250, HLA-A*2603, HLA-A*6601, HLA-A*6802, HLA-A*6901, HLA-B*1509, HLA-B*2720, HLA-B*3901, HLA-B*4801, HLA-C*0501, HLA-C*0602, HLA-C*0802, HLA-C*1203 and HLA-C*1402 had at least one CD8 T cell epitope containing the D614 amino acid (Table 4). Unlike the L5 and G1124 containing epitopes, D614 containing epitopes could start at several different positions including 606, 607, 610, 611, 612 and 614 with the epitope of YQDVNCTEV most frequently observed (Table 4).

**Table 4.**
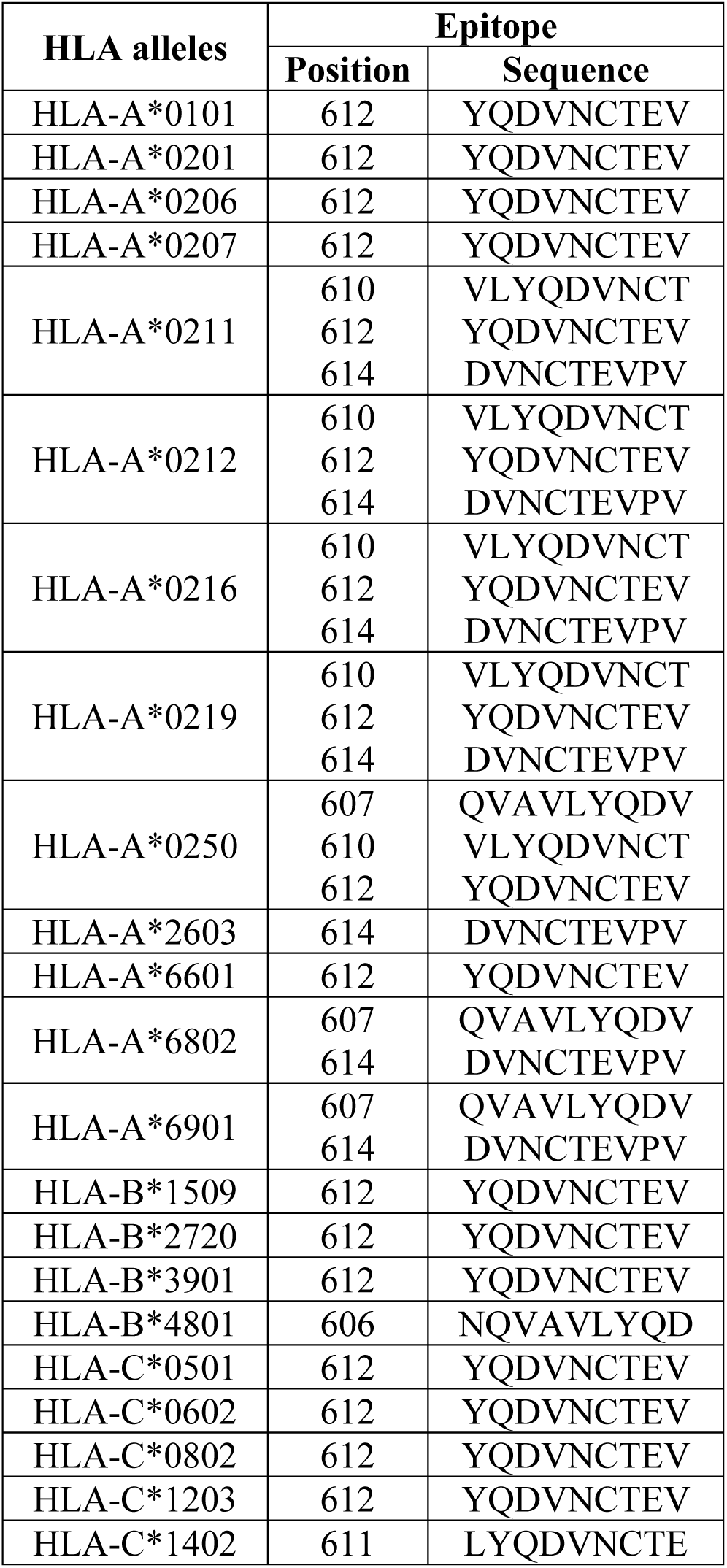
List of CD8 T cell epitopes containing the amino acid D614 on the reference S protein of SARS-CoV-2

Similarly, we reanalyzed the 9-mer CD8 T cell epitope for the reference S protein with a single D614G mutation. The resulting epitopes of each allele were compared to the corresponding epitopes derived from original reference S protein (Supplementary Table 2). The comparison data was summarized in Table 5. With a single D to G switch, there were seven HLA alleles including HLA-A*0203, HLA-A*0205, HLA-A*0206, HLA-A*2403, HLA-A*2501, HLA-A*2601 and HLA-C*1203 that obtained at least one CD8 epitope containing the replaced amino acid G614. Interestingly, nine HLA alleles including HLA-A*0101, HLA-A*0207, HLA-A*6802, HLA-A*6901, HLA-B*1509, HLA-B*3901, HLA-C*0501, HLA-C*0802 and HLA-C*1203 lost one CD8 epitopes each that were predicted using unmodified reference S protein sequence (Table 5). In addition to the direct gain or loss of CD8 T cell epitopes, several HLA alleles had their epitope binding affinities changed. These include the G614 containing epitopes with increased affinity for HLA-A*0211, HLA-A*0212, HLA-A*0216, HLA-A*0219, HLA-A*0250, HLA-A*6601, HLA-A*6802, HLA-A*6901, HLA-B*2720 and HLA-C*1402 (Table 5). Finally, we also observed there were other G614 containing epitopes with decreased affinity for alleles including HLA-A*0201, HLA-A*0206, HLA-A*0211, HLA-A*0212, HLA-A*0216, HLA-B*4801, HLA-C*0602 (Table 5). These results illustrated differential effects of D614G mutation on the capability to present S protein specific CD8 T cell epitopes by different HLA alleles.

**Table 5.**
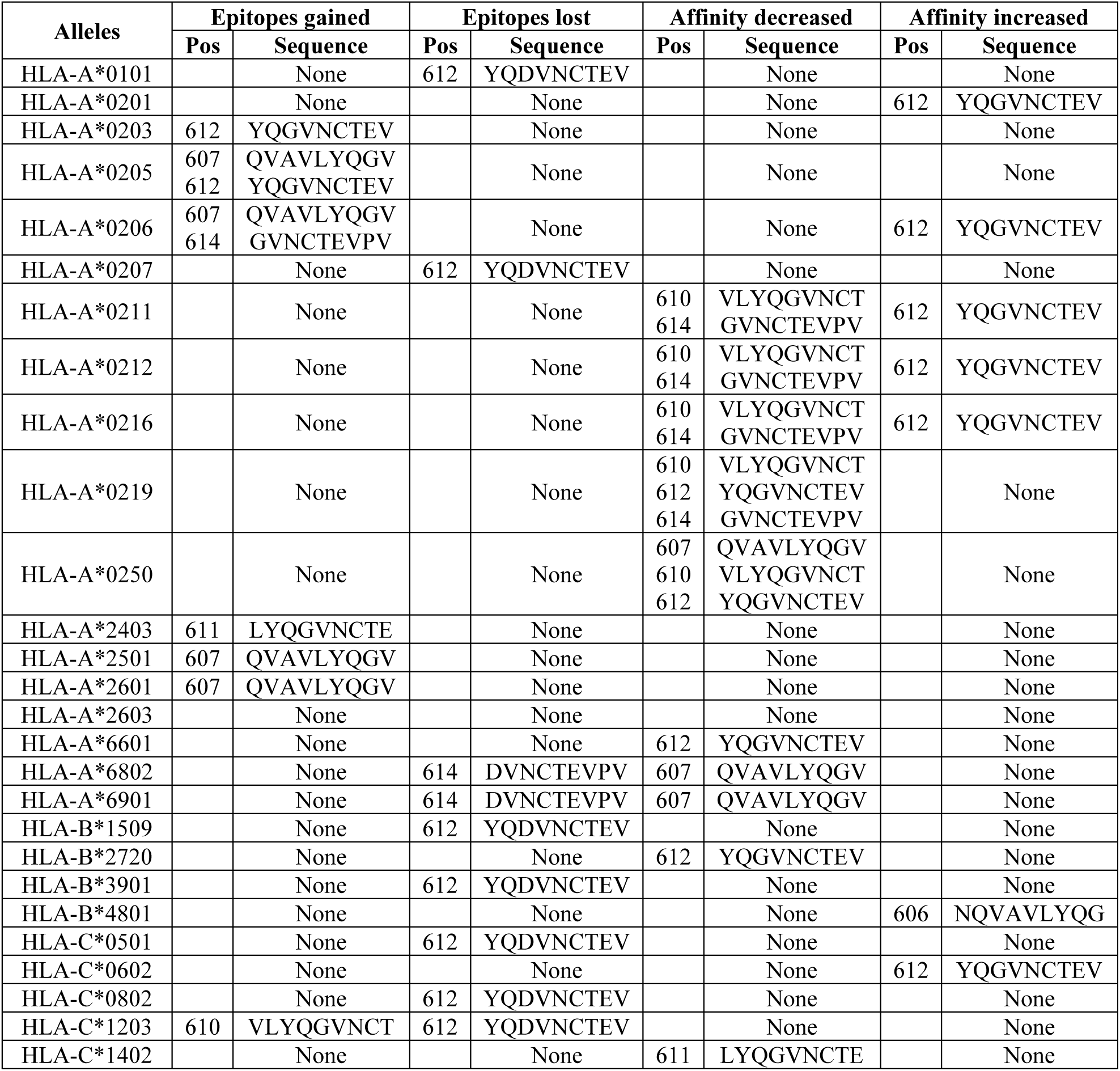
Effect of D614G mutation on the generation of SARS-CoV-2 S protein specific CD8 T cell epitopes by different HLA alleles

### Lack of seasonal CoVs cross-reactive CD8 T cell immunity to promote L5F, D614G and G1124V mutations

Currently, the detailed mechanisms driving these mutations on the SARS-CoV-2 S protein are not known. One possibility is that existing cross-reactive CD8 T cell immunity elicited by human seasonal CoVs could promote the virus to mutate and escape immune recognition in general population that are frequently targeted by seasonal CoVs. To test this, we went ahead to evaluate if there are shared CD8 T cell epitopes containing L5, D614 and G1124 from four representative seasonal CoVs(1,4). However, our pairwise alignment of the reference S protein with each of the four seasonal CoVs showed very low percent of identities (27% for HCoV-NL63, 31% for HCoV-HKU1, 28% for HCoV-229E and 33% for HCoV-OC-43). Further, multiple alignment of theses S protein sequences failed to identify any identical or similar CD8 T cell epitope motif to these we described above that include: FVFLVLLPL, FVSVNCDVV and several epitopes containing D614 (Table 4 and Fig 2). From these results, it was concluded that there were unlikely any cross-reactive CD8 T cells induced by seasonal CoVs that could promote the key mutations we observed.

**Fig 2.**
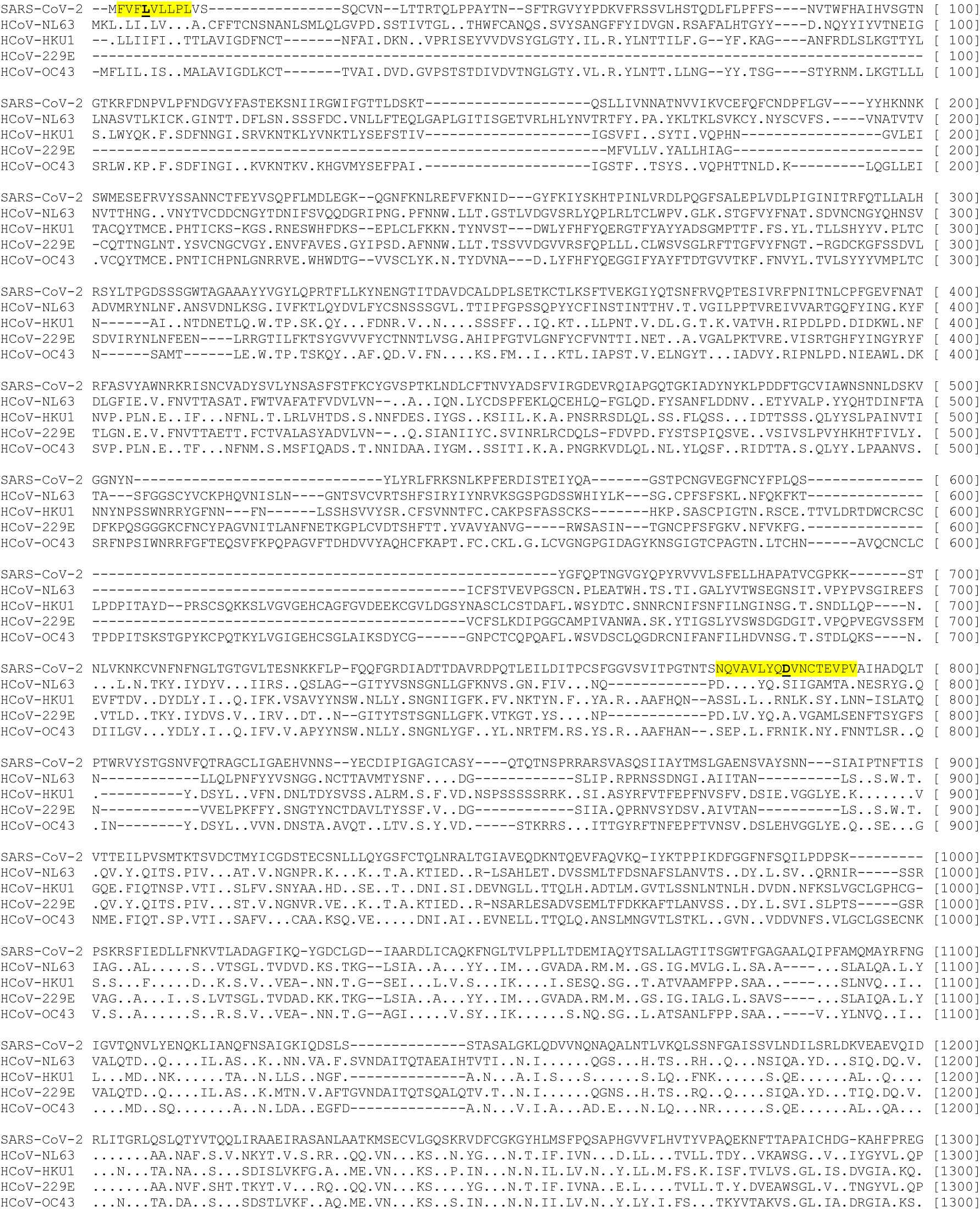
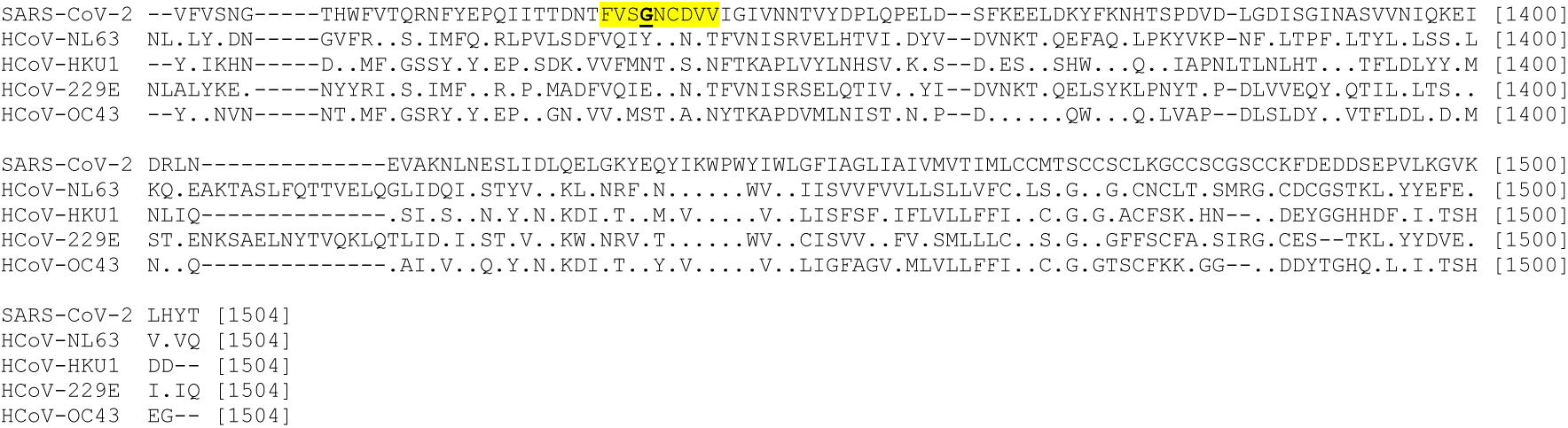
Multiple alignment of reference S protein of SARS-CoV-2 with four seasonal human CoVs (NL63, HKU1, 229E and OC43). The alignment was performed using Mega-X as described. The predicted CD8 T cell epitopes were highlighted in yellow and the corresponding mutational targets of L5, D614 and F1124 were bolded and underlined on the SARS-CoV-2. The accession number for S proteins of SARS-CoV-2, HCoV-NL63, HCoV-HKU1, HCoV-229E and HCoV-OC43 are QHD43416.1, APF29063.1, BBA20983.1, AAG48592.1 and CAA83661.1, respectfully.

## Discussion

CD8 T cell response specific for S protein of SARS-CoV has been well characterized, yet most of the data has primarily focused on HLA-A*0201(18,34–37). The epitopes of CD8 T cells and their specific responses against SARS-CoV in population with other HLA alleles are scarcely investigated. During SARS-CoV-2 infection, reduction of CD8 T cells is associated with worse prognosis and systemic inflammation(38). Yet, the specificities of these CD8 T cell and associated HLA genetic background were not provided.

In this study, we intended to provide a broad representation of S protein specific CD8 T cell epitopes of 80 different human HLAs for better understanding anti-SARS-CoV2 T cell immunity. Our results indicated a potential differential anti-S protein CD8 T cell response during COVID-19 infection in individuals with unique HLA alleles as the capacity of these alleles to present potential epitopes varies dramatically (Table 1). Among them, HLA-B*2705 is only capable of presenting 16 epitopes. This allele is highly associated with various forms of arthritis(39). It’s known that SARS-CoV-2 infection in aged population and people with underlying health conditions including immune disorders tend to be much severe and lethal(40). Additionally, during acute influenza infection and chronic HIV progression, viral escape mutations could occur on HLA-B*2705 restricted CD8 T cell epitopes(41,42). Although we didn’t find evidence that seasonal CoV-2 could induce cross-reactive CD8 T cells toward the three key mutations we identified, other mechanism may promote SARS-CoV-2 to mutate and escape HLA-B*2705 restricted CD8 T cell immunity. Population with Caucasian origins in several countries including Spain, Belgium, Austria, Netherlands and USA have from about 6% to 14% high frequency for this allele according to the allele frequency database (http://www.allelefrequencies.net/). Some of these countries have been experienced substantially high COVID-19 case numbers and/or mortalities. Therefore, SARS-CoV-2 infection rate, disease severity and immune responses in HLA-B*2705 individuals warrant thorough investigations.

In addition to HLA-B*2705, B44 alleles also had very low number of epitopes (17 for B4402 and 19 for B4403, Table 1). A recent epidemiologic study reported that HLA-B*44 and HLA-C*01 positive individuals were more susceptible to SARS-CoV-2 infections in Italy when compared with HLA-A*25, B*08, B*15:01, B*51, B*14, B*18, B*49, and C*03(23). Although we weren’t able to predict the epitopes for C*01 allele as it hasn’t been enlisted in NetMHC 4.0, each of A*25, B*08, B*15:01, B*51, B*14, B*18 and B*49 alleles we analyzed showed higher numbers of CD8 T cell epitopes than B*44 (Table 1), indicating potential broad anti-S protein CD8 T cell responses may provide better protection against SARS-CoV-2 infection.

Our data lacks experimental support, however, some of the S protein specific CD8 T cell epitopes presented in this study have been validated in SARS-CoV infection and utilized for monitoring human anti-SARS specific CD8 T cell responses using immunological techniques such as ELISPOT and tetramer staining(18,34–37). One of these epitopes is FIAGLIAIV that was identified in HLA-A*0201 patient infected with SARS-CoV(34). This epitope was predicted not only for HLA-A*0201 on SARS-CoV2 S protein, but also a total of other 21 alleles analyzed in this report including HLA-A*0202, HLA-A*0203, HLA-A*0205, HLA-A*0206, HLA-A*0207, HLA-A*0211, HLA-A*0212, HLA-A*0216, HLA-A*0217, HLA-A*0219, HLA-A*0250, HLA-A*2501, HLA-A*2601, HLA-A*2602, HLA-A*6802, HLA-A*6901, HLA-B*3901, HLA-B*4601, HLA-C*1203, HLA-C*0802, and HLA-C*1502 (Supplementary Table 2). The identified epitope RLNEVAKNL on SARS-CoV(37) was shared with 8 HLA-A, 4 HLA-B and 3 HLA-C alleles for SARS-CoV-2 in our study. Another SARS-CoV S protein specific CD8 T cell epitope, VLNDILSRL(36) were predicted on the SARS-CoV-2 S protein for 9 HLA-A alleles, 1 HLA-B and 3 HLA-C alleles. The epitope NLNESLIDL we identified for 10 HLA-A alleles was also characterized as a valid HLA-A*0201 restricted SARS-CoV CD8 T cell epitope in a published study(35).

Although recent studies agreed that D614G mutation could enhance SARS-CoV-2 infectivity and promote its transmission(27,43), regarding its effect on virulence, one study reported that G614 virus infection was associated with higher mortality(44), while the other study concluded no obvious effect on disease severity(27). One potential explanation for these disparate clinical findings is that the prevalent HLAs of the infected subjects in the two studies differ, which leads to divergent anti-S protein CD8 T cell responses, either toward the epitopes containing the G614 mutation alone and/or in combination with other epitopes we identified, as D614G mutation can occur simultaneously with other mutations on the S protein (Supplementary Fig 1 and data not shown). This is because first, our prediction result showed only about 25% of the total HLA alleles we analyzed could mount CD8 T cell responses targeting the epitopes containing D614 (Table 4). Secondly, among these HLA alleles that could generate D614 containing CD8 T cell epitopes, the epitopes they recognize and present to TCR differ (Table 5), which may lead to different TCR clonotypes and anti-viral efficacy(45,46). Third, our mutational analysis on a panel of CD8 T cell epitopes that have G614 also suggested possible different control outcomes by different HLA alleles as their bindings to the mutated epitope could be altered differently (Table 5). We believe pairing HLA typing with SARS-CoV-2 sequencing and testing anti-S protein CD8 T cell responses could allow precise assessment of clinical outcome of D614/G614 virus infection on individuals with different HLA alleles.

Our data of mutational effect of L5F suggests that the evolution of SARS-CoV2 virus targeting L5 might be eventually unsuccessful as the mutated epitope could enhance CD8 T cell recognition and killing through the enhanced interaction with most of the HLA alleles (Table 2). In contrast, the G1124V mutation appears to favor the virus to escape immune recognition as this mutation reduces epitope binding affinity of HLA alleles to a level that some are no longer able to bind (Table 3). However, the V1124 variant has not become as dominant as the G614. This could be due to other viral and host factors, such as viral structure stability, peptide-MHC stability and innate responses that prevent this variant to stand out further. Since the pandemic SARS-CoV-2 outbreak in several major countries hasn’t yet been controlled, the continual monitoring of these mutations on S protein are still necessary.

Based on the data and discussions provided here, we would like to encourage the research labs to carry out further validation and characterization of these candidate S protein epitopes and study their role in protecting SARS-CoV-2 infection and vaccine immunity. These epitopes should include not only the ones that had been identified for the S protein of SARS-CoV, but also uncharacterized epitopes such as these containing L/F5, D/G614 and G/V1124. We also recommend the clinical labs to organize and utilize resources to combine SARS-CoV-2 sampling and viral genome sequencing with HLA typing and S protein specific CD8 T cell immune testing to gather more useful data for better identifying risk groups and implementing policies that are suited for different geographical locations, resulting in more effective transmission control. Ultimately, we believe these efforts will provide more solid data for effective vaccine development and elimination of SARS-CoV-2 from human population.

## Material & Methods

### S protein sequence of reference virus

Full-length S protein amino acid sequence (accession number QHD43416.1) of the reference SARS-CoV-2 Wuhan-Hu-1 isolate (accession number MN908947) was downloaded in FASTA format from the NCBI GenBank.

### CD8 T cell epitope prediction

CD8 T cell epitope prediction was performed on NetMHC4.0 Web Server http://www.cbs.dtu.dk/services/NetMHC/ that utilizes artificial neural networks (ANNs)(47). A total of 36 HLA-A, 34 HLA-B and 10 HLA-C alleles (Table 1) was used for prediction. 9-mer peptides (epitopes) with rank score ≤2.0% were selected as positive HLA-binder. As the output format for the epitopes derived from NetMHC4.0 begins with 0, one was added to the resulting positions of all epitopes presented in this study to match the S protein sequence numbering.

### Mutational analysis

To identify mutations on SARS-CoV-2 S protein, we selected a total of 1000 complete or near-complete S protein sequences of from viruses isolated from countries in North America, Europe, Asia, Oceania, Africa and South America. These sequences were deposited in the NCBI database with collection dates ranging from January to June 2020 and showed at least 95% of query coverage for the reference S protein (accession number QHD43416.1). The list of the accession numbers for these sequences was provided in Supplementary table 1. Multiple alignment of these 1000 protein sequences against the reference S protein was conducted using NCBI BLASTP program under default conditions.

### Comparison of CoV S protein sequences

The S protein sequences of four common seasonal CoVs: HCoV-229E (AAG48592.1), HCoV-OC43 (CAA83661.1), HCoV-NL63 (APF29063.1), and HCoV-HKU1 (BBA20983.1) were used for comparison with the reference S protein of SARS-CoV 2 by multiple alignment in MEGA-X using MUSCLE under default conditions.

## Authors’ Contributions

EG initiated this study. EG and HG performed immune informatics and sequence analysis. EG and HG wrote and revised the manuscript. All authors read and approved the final manuscript.

## Acknowledgement

The health care professionals, researchers and laboratories around the world who have helped with collecting samples, sequencing and depositing the data for SARS-CoV-2 into NCBI database are sincerely appreciated.

This research received no financial support from any funding agencies.

